# Change in terrestrial human footprint drives continued loss of intact ecosystems

**DOI:** 10.1101/2020.05.04.077818

**Authors:** Brooke A. Williams, Oscar Venter, James R. Allan, Scott C. Atkinson, Jose A. Rehbein, Michelle S. Ward, Moreno Di Marco, Hedley S. Grantham, Jamison Ervin, Scott Goetz, Andrew J. Hansen, Patrick Jantz, Rajeev Pillay, Susana Rodríguez-Buriticá, Christina Supples, Anne L. S. Virnig, James E.M. Watson

## Abstract

Our ability to map humanity’s influence across Earth has evolved, thanks to powerful computing, a network of earth observing satellites, and new bottom-up census and crowd-sourced data. Here, we provide the latest temporally inter-comparable maps of the terrestrial Human Footprint, and assessment of change in human pressure at global, biome, and ecoregional scales. In 2013, 42% of terrestrial Earth could be considered relatively free of anthropogenic disturbance, and 25% could be classed as ‘wilderness’ (the least degraded end of the human footprint spectrum). Between 2000 and 2013, 1.9 million km^2^ - an area the size of Mexico - of land relatively free of human disturbance became highly modified. The majority of this occurred within tropical and subtropical grasslands, savannah, and shrubland ecosystems, but the rainforests of Southeast Asia also underwent rapid modification. Our results show that humanity’s footprint is eroding Earth’s last intact ecosystems, and greater efforts are urgently needed to retain them.

## Introduction

Humans have influenced the terrestrial biosphere for millennia, converting much of Earth’s surface to anthropogenic land uses^1^. Nevertheless, there are still some ecosystems that remain free from significant human pressure, thereby providing crucial habitats for imperilled species^2,3^ and maintaining the ecosystem processes that underpin planetary life-support systems^4,5^. As a consequence, calls for the global identification, monitoring, and retention of the remaining lands that are relatively free of direct anthropogenic disturbance are increasing^6–8^.

Over the past two decades, cumulative pressure maps that combine remotely-sensed data with survey data are being increasingly used to assess the full range of human pressures on land spatially^9^. These advances have facilitated the mapping of Earth’s remaining marine and terrestrial wilderness^8,10,11^, improved measures and estimates of species extinction risk^12^, underpinned broader assessments of human impacts on ecosystems13 and biodiversity^14–16^, and enabled the identification of protected areas and world heritage sites in danger14,17,18. The results of these mapping efforts are influencing global policy discussions^6,19^, and informing on-the-ground decisions about where to undertake biodiversity conservation action^20–22^.

Here, we provide the latest global maps of cumulative human pressure^23,24^ for the years 2000, 2005, 2010, and 2013, and use them to assess how changes in human pressure are altering Earth’s terrestrial ecosystems. We used a human footprint threshold of <4 (on 0 – 50 scale) to identify where land is considered ecologically intact (below the threshold) or highly modified and thus ecologically degraded (equal to or above the threshold). Areas below this threshold are ecosystems that may be subject to some level of human pressure (for example low-density transitory human populations or pasture lands grazed at a low intensity), but still contain the majority of their natural habitat and ecological processes^14,25^. This threshold has been found to be robust from a species conservation perspective because once surpassed, species extinction risk increases dramatically^12^, and several ecosystem processes are altered^12,16,26^.

We assess transitions from intact to highly modified land at global, biome, and ecoregional scales^27^ and ascertain which nations contain Earth’s remaining intact systems, and had the greatest amounts of habitat loss. Previous global assessments of human pressure have attempted to identify at risk ecosystems by determining a ‘safe limit’ of biodiversity loss for ecosystem functionality^28,29^, assessing protection levels^30^, and analysing habitat conversion using land cover^31,32^. But all of these ignore a broad range of threats that occur beyond land use such as accessibility via roads, railways and navigable waterways, human population density, and light pollution. These pressures have environmental impacts well beyond the local development footprint^33,34,36^. As such, our results provide the latest spatially explicit understanding of the state of human pressure on the natural environment, and how it is changing over time. We show that the human footprint methodology can be continually updated and, when more recent data becomes available, allow for assessment of habitat loss at scales relevant to planning activities.

## Results

### State of terrestrial Earth

As of 2013, 55.8 million km^2^ (41.6%) of Earth’s surface was intact (which includes wilderness, human footprint of <4), and 33.5 million km^2^ (25.0%) was wilderness (human footprint of <1). The remaining (human footprint of ≥4) 78.4 million km^2^ (58.4%) was under moderate or intense human pressure (and therefore highly modified), which was widespread, encompassing over half the area of 11 (or 78.6%) of Earth’s 14 biomes (Figure 1). Temperate broadleaf and mixed forests were the most altered biome, with 11.6 million km^2^ (91.0%) being highly modified, followed by tropical and subtropical dry broadleaf forests with 2.72 million km^2^ (90.5%), and Mediterranean forests, woodlands and scrubs with 2.88 million km^2^ (89.7%). Wilderness areas have all but disappeared in many biomes, for example, only 82,000 km^2^ (0.81%) remained in temperate grasslands, savannahs, and shrublands, 29,000 km^2^ (0.96%) in tropical and subtropical dry broadleaf forests, and just 12,000 km^2^ (1.69%) in tropical and subtropical coniferous forests.

**Figure 1.**
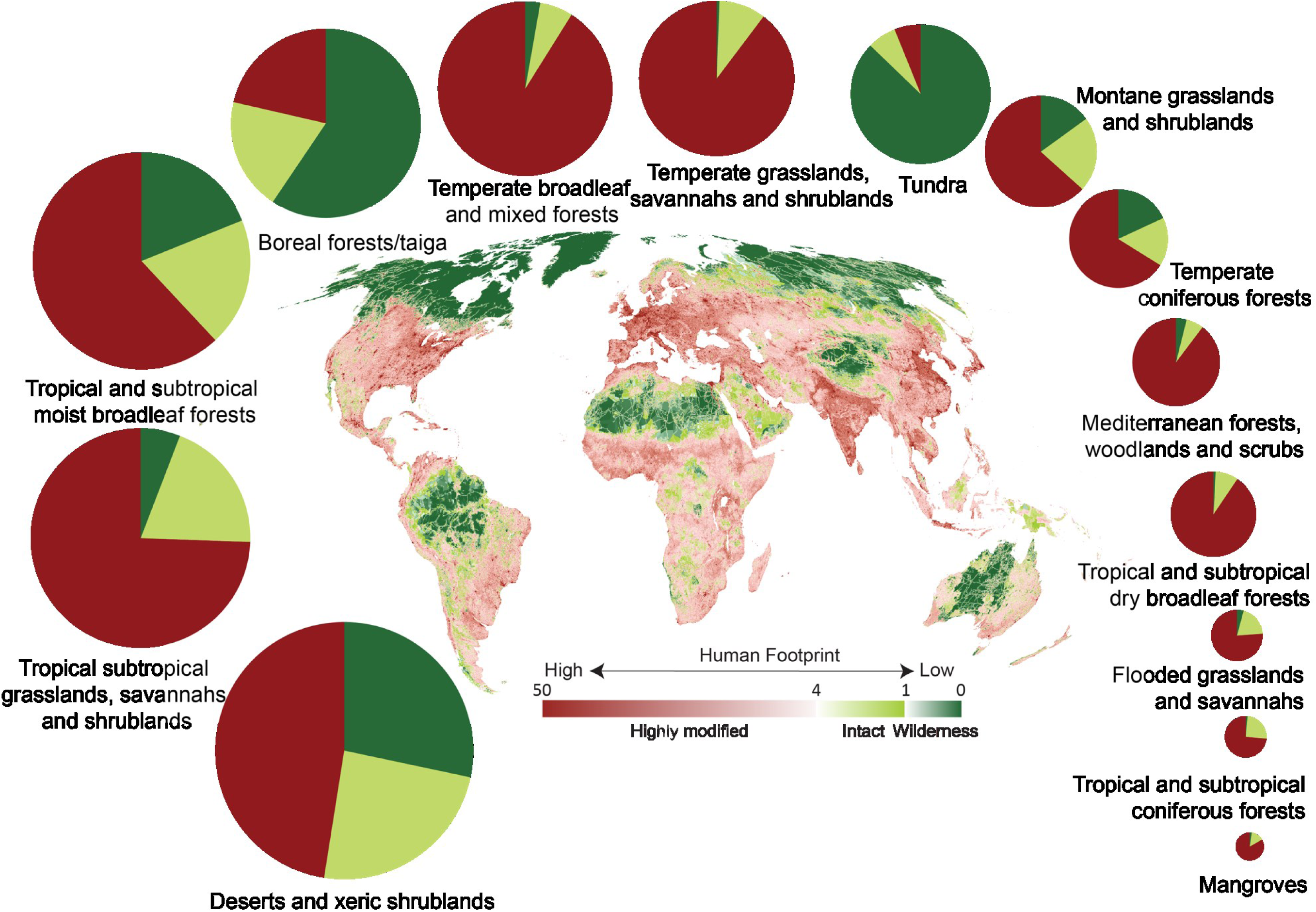
The global human footprint map for the year 2013. The surrounding pie charts represent the proportion of each terrestrial biome that was completely free of direct anthropogenic disturbance (‘wilderness’, dark green, human footprint value of <1), relatively free of direct anthropogenic disturbance (‘intact’, light green, human footprint value of <4 and ≥1), or highly impacted by anthropogenic disturbance (‘highly modified’, red, human footprint value of ≥4) in the year 2013. Circles sizes represent relative biome area.

Earth’s 14 biomes consist of 795 ecoregions, which represent distinct biotic assemblages and abiotic features (such as landforms) at a finer scale than biomes^27^. We found the entire extent of 46 (5.76%) ecoregions were highly modified. These 46 ecoregions span 10 biomes, with the majority located in tropical and subtropical moist broadleaf forests (n = 17, 37.0%), tropical and subtropical dry broadleaf forests (n=6, 13.0%), and temperate broadleaf and mixed forests (n=6, 13.0%). One-quarter of all ecoregions (n=187) have lost all wilderness.

The majority of land in tundra, boreal and taiga forests, and deserts and xeric shrubland biomes remains intact. At the ecoregion level, just 52 (6.53%) still have >90% of their land intact, and a mere 21 (2.64%) are >90% wilderness. These ecoregions with >90% wilderness are found in just four biomes, tundra (n = 12), boreal forests/taiga (n = 5), tropical and subtropical moist broadleaf forests (Rio Negro campinarana and Juruá-Purus moist forests), and tropical and subtropical grasslands, savannahs and shrublands (Northwestern Hawaii scrub).

### Contemporary changes in human pressure

Between 2000 and 2013, 25.4 million km^2^ (18.9%) of Earth’s terrestrial surface deteriorated (human pressure increased), while only 8 million km^2^ (5.96%) improved (human pressure decreased; Figure 2). This increase in human pressure was substantial across 1.89 million km^2^ of Earth’s intact lands, an area the size of Mexico, that these places can be classified as highly modified (i.e. they transitioned from below to above the human footprint threshold of 4; Figure 3). During the same time period, over 1.1 million km^2^ of wilderness was lost (human footprint increasing above 1), with 67,000 km^2^ of that wilderness becoming highly modified (human footprint increasing from below 1 to above 4; Figure 2; Figure 3).

**Figure 2.**
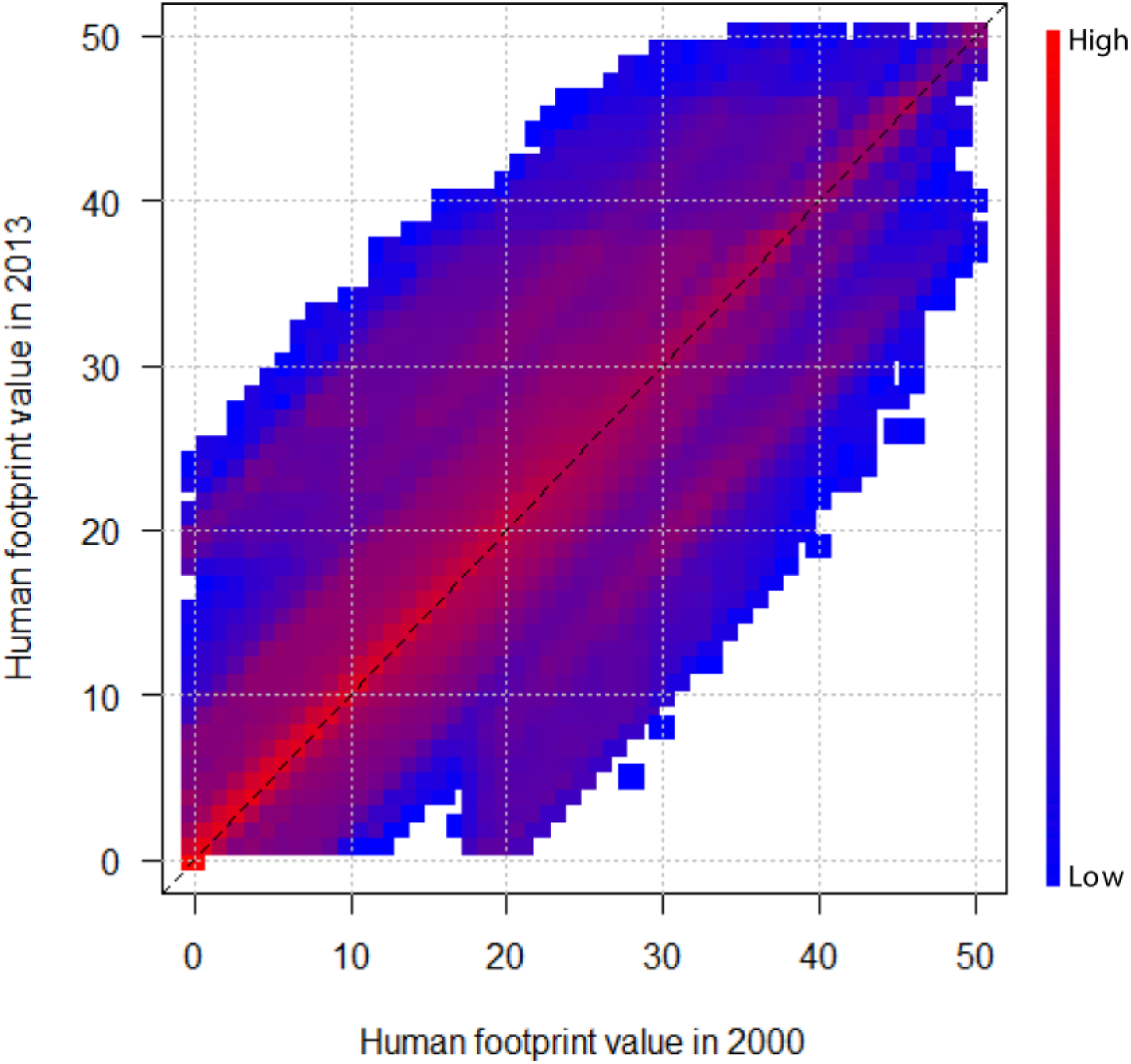
Density plot depicting change in the global terrestrial human footprint between the years 2000 and 2013 (n = 134,154,306). The x-axis represents the human footprint value of a pixel in the year 2000, and the y-axis represents the human footprint value of that pixel in the year 2013. The number of pixels that made that particular transition are represented by the colour within the plot. Red represents a high number of pixels and blue represents low. Legend is log-scaled. Between 2000 and 2013, 25,348,514 km^2^ (18.9%) of pixels deteriorated (human pressure increased), while 7,995,464 km^2^ (5.96%) improved (human pressure decreased).

**Figure 3.**
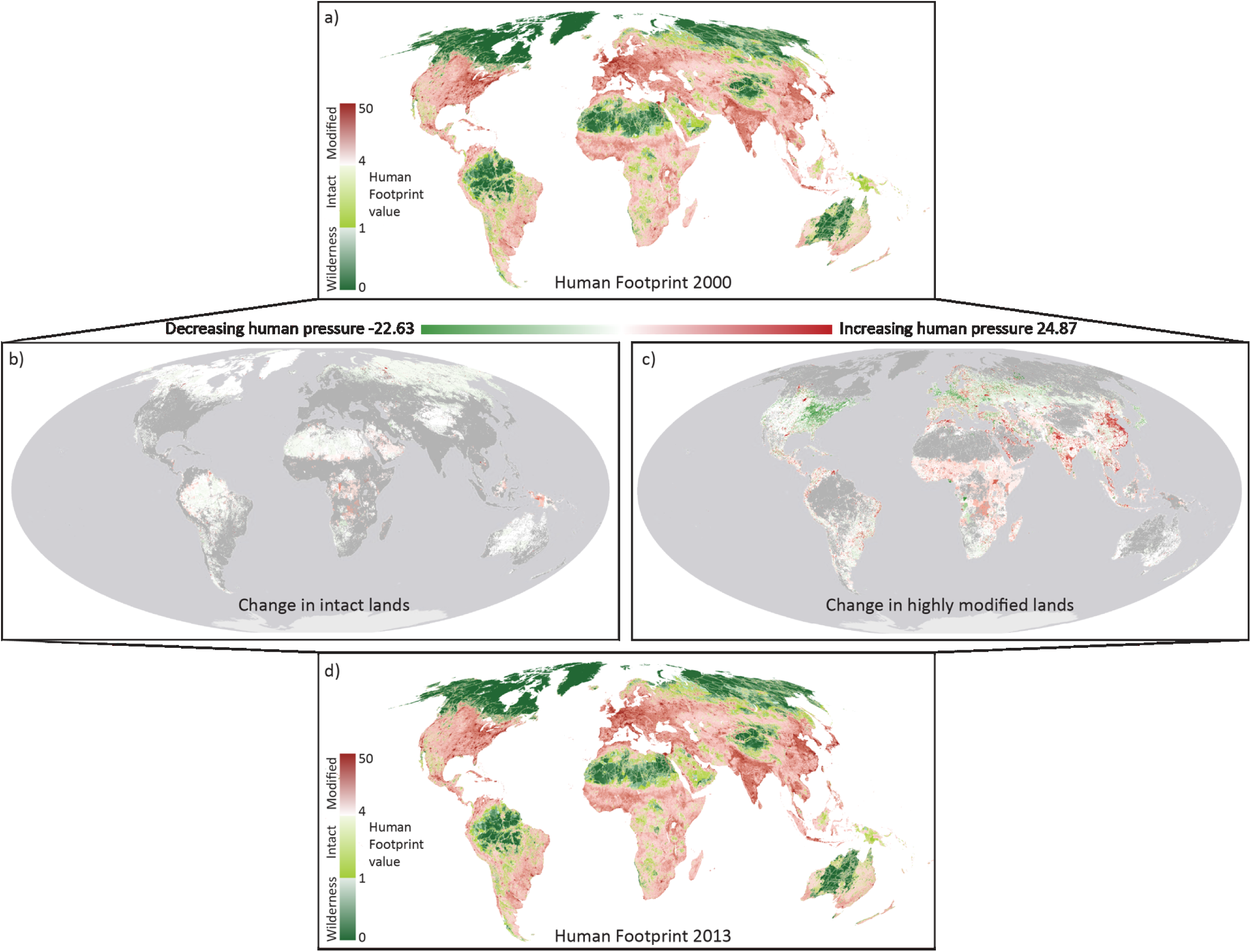
The global human footprint map for the year 2000 (a). Areas completely free of direct anthropogenic disturbance (‘wilderness’, dark green, human footprint value of <1), relatively free of direct anthropogenic disturbance (‘intact’, light green, human footprint value of <4 and ≥1), or highly impacted by anthropogenic disturbance (‘highly modified’, red, human footprint value of ≥4). The change between 2000 and 2013 within each 2000 state can be seen for intact land (b) and highly modified land (c), which leads to the 2013 state (d).

Intact lands were lost in all biomes during the assessment period, with the highest loss occurring in tropical and subtropical grassland, savannah and shrublands (655,000 km^2^ was lost representing 11.3% of all intact lands within the biome, an area approximately the size of France; Figure 4). The tropical and subtropical moist broadleaf forests and mangrove biomes also lost substantial areas of intact land (559,000 km^2^, 6.90% and 9,000 km^2^, 14.7% respectively). While the largest absolute loss of intact lands occurred in savannah and woodland ecoregions, the largest proportional losses occurred in tropical forest ecoregion types. For example, intact areas were completely lost in seven forested ecoregions including the Louisiade Archipelago rainforests (Papua New Guinea), and Sumatran freshwater swamp forests (Indonesia; see Supplemental 1).

**Figure 4.**
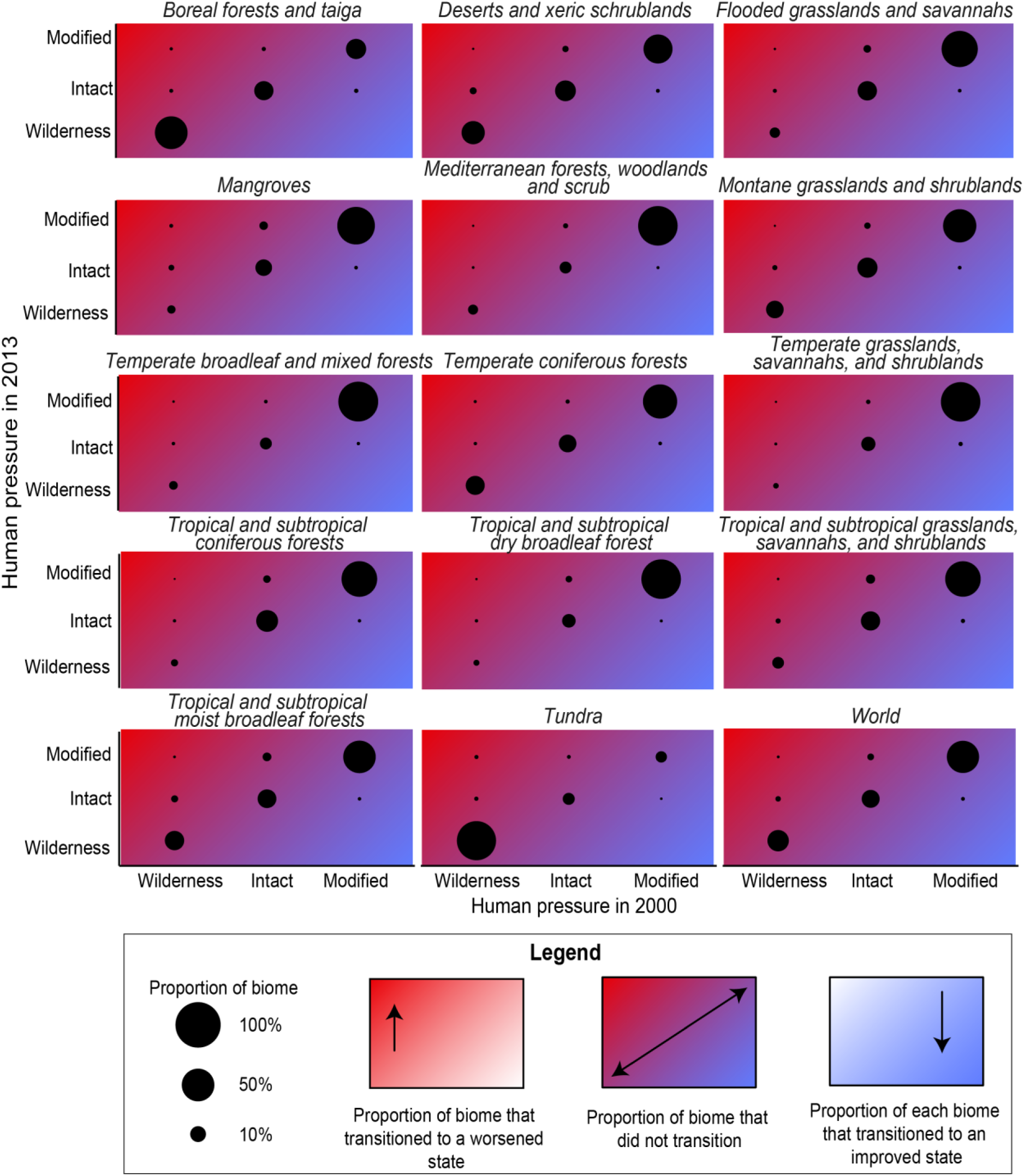
The proportion of biome that transitioned between wilderness (human footprint value of <1), intact (human footprint value between <4 and ≥1), and highly modified (human footprint value of ≥4) states between 2000 and 2013, represented by circles. If part of the biome transitions to a worsened condition it moves upwards into the red area, if part of the biome does not transition it remains on the diagonal, and if part of the biome improves, it moves downwards into the blue area. For exact values see Supplemental 1.

The largest losses of wilderness between 2000 and 2013 occurred in biomes that contained the largest areas of wilderness in 2000. For example, deserts and xeric shrublands lost 426,000 km^2^ (5.08%) of their remaining wilderness. This was concentrated in desert, woodland and savannah ecoregion types (see Supplemental 1). Wilderness in the tundra and boreal/taiga forests suffered the most extreme transitions, with 22,000 km^2^ and 15,000 km^2^ respectively changing from wilderness to highly modified land (human footprint <1 to ≥ 4) (Figure 4). The ecoregions of the Russian tundra and taiga lost the most wilderness, for example, the Yamal-Gydan tundra lost 8,000 km^2^, and the East Siberian taiga lost 5,000 km^2^.

### National responsibility

In 2013, only 26 nations (out of 221) had the majority (>50%) of their land intact. Excluding island territories, the two countries with the highest proportion of intact land included Guyana (88.8% of country; 187,000 km^2^) and Suriname (88.5%; 125,000 km^2^). The African continent contained 11 ecoregions that lost the largest areas of intact land. Between 2000 and 2013, more intact land was lost in the Democratic Republic of the Congo (DRC) than any other country (316,000 km^2^; 13.6% of the country; 37.3% of its intact lands). This was followed by Indonesia and Brazil which lost 122,000 km^2^ (6.98% of the country or 20.2% of its intact lands) and 87,000 km^2^ (29% of the country or 1.88% of its intact lands) respectively.

Russia, Canada, Brazil, and Australia are responsible for the largest areas of Earth’s remaining intact areas (which includes wilderness, human footprint score of <4). Combined, these four countries harbour more than 60% of Earth’s wilderness (human footprint score of <1, Figure 5). Brazil also lost the most wilderness (human footprint increasing above 1) of any country (109,880 km^2^, 3.87% of its wilderness area). The largest areas of wilderness lost to high levels of human modification (human footprint increasing above 4) were in Russia (23,000 km^2^), Canada (10,000 km^2^), and Brazil (6,000 km^2^).

**Figure 5.**
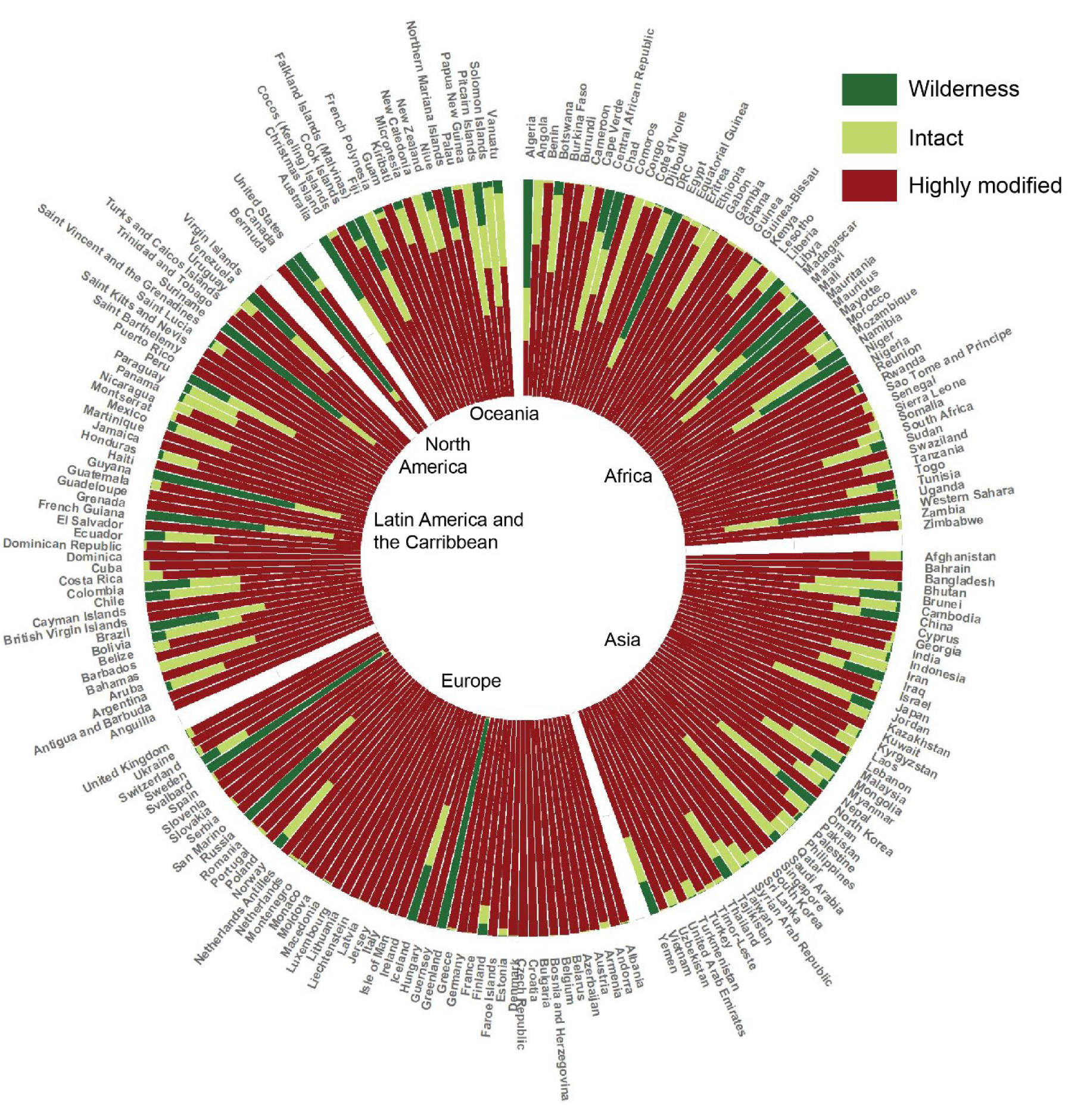
Proportion of each country’s terrestrial land that was completely free of direct anthropogenic disturbance (‘wilderness’, dark green, human footprint value of <1), relatively free of direct anthropogenic disturbance (‘intact’, light green, human footprint value between <4 and ≥1), or highly impacted by anthropogenic disturbance (‘highly modified’, red, human footprint value of ≥4) in the year 2013.

## Discussion

The terrestrial human footprint presented here is one of the most comprehensive and up-to-date measures of cumulative human pressure across Earth, and will be continuously improved as more data on the eight included pressures (built environments, population density, night-time lights, crop lands, pasture lands, accessibility via roads, railways, and navigable waterways) become available. While this latest update is already seven years out of date, advances in data generation and modelling^35^ will facilitate more rapid updates of the human footprint in the near future. Our analyses show that between 2000 and 2013 substantial areas of intact land, including wilderness areas, have been lost. This loss has profound implications for the biodiversity that require intact land for their continued survival^3^, and for people who rely on the services that intact ecosystems provide^8,37^. The transition from intact ecosystems to highly modified land is the greatest predictor of why species face increasing extinction risk^12^ as this transition is where habitat is considered functionally unavailable for many terrestrial vertebrates^38,39^. This transition also negatively impacts wildlife population viability, because intact ecosystems are proven strongholds for genetic diversity40. Climate change mitigation efforts are also undermined by these losses because intact lands make crucial contributions to the residual terrestrial carbon sink^37,41^. For example, a recent study found that carbon impacts of intact forest loss are 626% worse than originally estimated^41^.

We also demonstrate that patterns of degradation due to increasing human pressure are now changing within biomes. Past studies note that dry forested biomes have suffered the highest rates of habitat loss^25,31^ but our results now show that recent increases in human pressure predominantly occurred in tropical savannah and grassland ecosystems, which lost 11.3% of their intact area between 2000 and 2013. This finding is consistent with previous evidence that savannahs are the current development frontier in many regions worldwide^42,43^. Proactive conservation planning is urgently needed to prevent the last intact savannahs, such as Australia’s northern savannahs^44^ and Colombia’s Llanos in the Orinoquia region^43^, suffering the same losses that occurred in places such as Brazil’s Cerrado^45^. Conservation planning needs to utilise tools that take into account past and future risk, so that preventative conservation action can be implemented in places where development is most likely to occur^46–48^. Our analysis helps inform where proactive conservation planning activity must occur, and demonstrates the potential of human pressure mapping for informing global conservation action.

Nearly three decades ago, the world came together to ratify the Rio Conventions, including the Convention on Biological Diversity (CBD), the UN Convention to Combat Desertification (UNCCD), and the UN Framework Convention on Climate Change (UNFCCC). Despite the fact that almost all nations are signatories on these three international environmental agreements, intact habitats continue to be lost at a rapid rate^49^, including within the borders of many signatory nations, such as the DRC, Indonesia, and Brazil. One possible explanation for this trend is the challenge of collectively identifying intact landscapes and then using this information to take coordinated action across the globe to protect them. Given the growing body of scientific evidence demonstrating the exceptional value of intact ecosystems (including wilderness areas) for conserving biodiversity^50^, mitigating climate change^41^, and providing essential ecosystem services^37^, the importance of data on intactness should be elevated when undertaking efforts to develop international and national targets and shaping actions under these Conventions. For example, at the end of 2020 nations that are party to the Convention on Biological Diversity will sign off on the Post-2020 Global Biodiversity Framework that will set global targets on nature for the coming decades. Negotiations around the Post-2020 Framework present an opportunity for countries to include targets specifically for the protection and complete retention of intact ecosystems^8^.

Halting the loss of intact ecosystems cannot be achieved alongside current trajectories of development, population growth, and resource consumption^51^. Retention of Earth’s remaining intact lands can only be achieved through a combination of strategic policy mixes that better regulate deleterious activities across all sectors, levels of governance and jurisdictions, and on the ground site-based action such as well-resourced protected areas in conjunction with other effective area-based conservation measures (OECMS) such as payment schemes for safeguarding ecosystem services^51–54^. While many pathways on how intact retention can be achieved are being developed^8,51,54,55^, the challenge is ensuring action occurs at the scale and speed necessary to ensure all intact ecosystems are secured.

The highest losses of intact lands occurred in African nations, where the highest biodiversity impact from future socio-economic development is also predicted to occur56. Parts of Africa also have the largest gap between food consumption and production in the world, we can therefore only infer that increasing agricultural production is a key driver of savannah and grassland loss^57,58^. Other regions experiencing extreme levels of intact ecosystem loss are the rainforests of Indonesia (which covers 1.3% of Earth but contains 10% of the world’s plants, 12% of mammals, 16% of reptile–amphibians, and 17% of birds^59^) and Papua New Guinea (which covers less than 1% of Earth but contains 5% of its biodiversity^60^). This extreme habitat loss is likely due to the spike in habitat conversion to grow cash crops such as oil palm^61,62^, driven by international demand^63^. Thus, research must be oriented to understanding these drivers, and subsequently to find mechanisms that facilitate socio-economic development without further degrading intact ecosystems^51,64^.

## Conclusion

We have presented the latest comprehensive assessment of humanity’s footprint on terrestrial Earth using the best available data. We find human pressure is extending ever further into the last ecologically intact, and wilderness areas. With important policy discussions on the Convention on Biological Diversity’s Post-2020 Global Biodiversity Framework well underway ^65^, this is a timely opportunity for nations to take stock and to set explicit targets for retaining Earth’s remaining intact lands. Proactively protecting Earth’s intact ecosystems is humanity’s best mechanism for protecting against climate change, ensuring large-scale ecological and evolutionary processes persist, and safeguarding biological diversity into the future.

## Experimental procedures

### Overview

We updated the Human Footprint^23^ terrestrial cumulative human pressure maps for the years 2000, 2005, 2010 and 2013 and used it to define the state of Earth’s biomes, ecoregions and countries, and their transitions between states between 2000 and 2013. All analyses, and creation of the human footprint maps, were conducted in the Mollweide equal area projection at 1 km^2^ resolution.

### Updating the human footprint

To recreate the human footprint maps we followed broadly the methods developed by Sanderson and colleagues^24^ and Venter and colleagues^23^. Significant areas missing in the original Human Footprint^23^ (which carried over into subsequent releases), including Azerbaijan, areas along the western former-USSR border, and along the Orange River in South Africa, among others, have been included in this update. We used data on human pressures across the periods 2000 to 2013 to map: 1) the extent of built human environments, 2) population density, 3) electric infrastructure, 4) crop lands, 5) pasture lands, 6) roadways, 7) railways, and 8) navigable waterways. To facilitate comparison across pressures we placed each human pressure within a 0–10 scale, weighted within that range according to estimates of their relative levels of human pressure following Sanderson and colleagues^24^. The resulting standardized pressures were then summed together to create the standardized human footprint maps for all non-Antarctic land areas. Pressures are not intended to be mutually exclusive, and many will co-occur in the same location. Three pressures only had data from a single time period or have poorly annotated temporal information, and these are treated as static in the human footprint maps.

We used free and open-source GRASS GIS 7.2.2^66^ to create a series of scripts that integrate the spatial data on human pressures, yielding 134,064,303 pixels for Earth’s terrestrial surface (excluding Antarctica). For any grid cell, the human footprint can range between 0–50. We carried out a validation of the human footprint map using visual interpretation of high resolution imagery across 3114 × 1 km^2^ sample plots randomly located across the Earth’s non-Antarctic land areas. We found strong agreement between the human footprint measure of pressure and pressures scored by visual interpretation of high resolution imagery, with a root mean squared error (RMSE) of 0.116 and a Kappa statistic of 0.806 (P < 0.01). For further details on the validation exercise see Supplemental 2. The following sections (and Table S1) describe in detail the source data for each pressure, the processing steps applied, and the rationale behind the pressure weighting. The code and underlying data for generating these maps is available online at https://github.com/scabecks/humanfootprint_2000-2013, and can be used to easily regenerate them with updated or alternate datasets, as well as to apply the same methodology at national or regional scales.

### Built environments

Built environments, in the context of the human footprint, are anthropogenic areas that represent urban settings, including buildings, paved land and urban parks. These environments do not provide viable habitats for many species of conservation concern, nor do they provide high levels of ecosystem services^67–70^. As such, built environments were assigned a pressure score of 10.

To map built environments, we used the Defence Meteorological Satellite Program Operational Line Scanner (DMSP-OLS) composite images which gives the annual average brightness of 30 arc second (~1 km at the equator) pixels in units of digital numbers (DN)^71,72^. This data was collected from six different satellite missions over the period 1992 to 2013. We extracted data for the years 2000, 2005, 2010, and 2013, and all datasets were then inter-calibrated to facilitate comparison^71^. Using the DMSP-OLS datasets, we considered pixels to be ‘built’ if they exhibited a calibrated DN greater than 20. This threshold is based on a global analysis of the implications of a range of thresholds for mapped extent of cities^73^, and visual validation against Landsat imagery for 10 cities spread globally.

The DMSP-OLS has limitations for the purpose of mapping human settlements, including hyper sensitivity of the sensors causing detection of over-glow adjacent to built environments73 and bright lights associated with gas flaring from oil production facilities^74^. However, no other data exist to map built environments in a consistent way globally over our time horizon. While more recent satellite platforms launches – such as VIIRS – offer higher spatial resolution and greater light sensitivity^75^ than DMSP-OLS, they aren’t presently comparable or integrated across the temporal range we required.

### Population density

The intensity of human pressure on the environment is often associated with proximity to human populations, such as human disturbance, hunting and the persecution of non-desired species^76^. Even low-density human populations with limited technology and development can have significant impacts on biodiversity^77,78^.

We incorporated human population density using the Gridded Population of the World dataset developed by the Centre for International Earth Science Information Network (CIESEN)^79^. The dataset provides a 1 km^2^ gridded summary of population census data for the years 2000, 2005, 2010, and 2013. We used linearly interpolated densities for year 2013 from data for years 2010 and 2015. For all locations with more than 1000 people km^−2^, we assigned a pressure score of 10. For more sparsely populated areas with densities lower than 1000 people km^−2^, we logarithmically scaled the pressure score using,

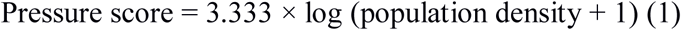

Human population density is scored in this way under the assumption that the pressures people induce on their local natural systems increase logarithmically with increasing population density, and saturate at a level of 1000 people km^−2^.

### Night-time lights

The high sensitivity of the DMSP-OLS^72^ dataset provides a means for mapping the sparser electric infrastructure typical of more rural and suburban areas. In 2009, 79% of the lights registered in the DMSP-OLS dataset had a Digital Number less than 20, and are therefore not included in our ‘built environments’ layers. However, these lower DN values are often important human infrastructures, such as rural housing or working landscapes, with associated pressures on natural environments.

To include these pressures, we used the inter-calibrated DMSP-OLS layers^71,72,80^ used for the built environments mapping. The 2013 calibration parameters were conveyed through personal communications from the creators of the dataset, and are not yet published. The equations for inter-calibrating across years are second order quadratics trained using data from Sicily, which was chosen as it had negligible infrastructure change over this period and where DN average roughly 14^72^. For our purposes, DN values of six or less where excluded from consideration prior to calibration of data, as the shape of the quadratic function leads to severe distortion of very low DN values. The inter-calibrated DN data from 2000 were then rescaled using an equal quantile approach into a 0–10 scale. To scale the data, we divided the calibrated night light data into 10 equal sample bins (each bin with a DN greater than 1 contains the same number of pixels) based on the DN values and then assigned them scores of 1 through 10, starting with the lowest DN bin. DN values of 0 were assigned a score of 0. The thresholds used to bin the 2000 data where then used to convert the 2005, 2010, and 2013 data into a comparable 0–10 scale.

### Crop and pasture lands

Crop lands vary in their structure from intensely managed monocultures receiving high inputs of pesticides and fertilizers, to mosaic agricultures such as slash and burn methods that can support intermediate levels of natural values^82,84^. For the purposes of the human footprint, we focused only on intensive agriculture because of its greater direct pressure on the environment, as well as to circumvent the shortcomings of using remotely sensed data to map mosaic agriculture globally, namely the tendency to confound agriculture mosaics with natural woodland and savannah ecosystems^86^.

Spatial data on remotely sensed agriculture extent were extracted from the MERIS CCI Landcover annual dataset ^81^. Although intensive agriculture often results in whole-scale ecosystem conversion, we gave it a pressure score of 7, which is lower than built environments because of their less impervious cover.

Pasture lands cover 22% of the Earth’s land base or almost twice that of agricultural crop^83^, making them the most extensive direct human pressure on the environment. Land grazed by domesticated herbivores is often degraded through a combination of fencing, intensive browsing, soil compaction, invasive grasses and other species, and altered fire regimes^88^. We mapped grazing lands for the year 2000 using a spatial dataset that combines agricultural census data with satellite derived land cover to map pasture extent^83^. We assigned pasture a pressure score of 4, which was then scaled from 0–4 using the percent pasture for each 1 km^2^ pixel.

### Roads and railways

As one of humanity’s most prolific linear infrastructures, roads are an important direct driver of habitat conversion^89^. Beyond simply reducing the extent of suitable habitat, roads can act as population sinks for many species through traffic induced mortality^90^. Roads also fragment otherwise contiguous blocks of habitat, and create edge effects such as reduced humidity^91^ and increased fire frequency that reach well beyond the roads immediate footprint^92^. Finally, roads provide conduits for humans to access nature, bringing hunters and nature users into otherwise wilderness locations^93^.

Data from OpenStreetMaps (OSM) on roads and railways was extracted from the global OSM planet database^85^. We include all categories of tagged highway in the OSM planet database. OSM is a volunteer driven, open-source global mapping project that has grown enormously in spatial completeness since its inception in 2004^94^. The volume and coverage of global transportation networks in the OSM database has far surpassed previously available roads data (e.g., gRoads^95^) which was used in earlier iterations of the Human Footprint^23^; however, the OSM dataset still does not provide full coverage outside of urban areas in some global regions, notably in central Africa, at the time of data extraction. Therefore, to benefit both from the larger OSM database while maintaining road coverages in regions that are currently poorly mapped in OSM, we merged the OSM data with gRoads data. The merged dataset performed best globally when we validated the three data layers (gRoads only, OSM only, and the union of gRoads/OSM).

We mapped the direct and indirect influence of roads by assigning a pressure score of 8 for 0.5 km out for either side of roads, and access pressures were awarded a score of 4 at 0.5 km and decaying exponentially out to 15 km either side of the road. While railways are an important component of our global transport system, their pressure on the environment differs in nature from that of our road networks. By modifying a linear swath of habitat, railways exert direct pressure where they are constructed, similar to roads. However, as passengers seldom disembark from trains in places other than rail stations, railways do not provide a means of accessing the natural environments along their borders. The direct pressure of railways where assigned a pressure score of 8 for a distance of 0.5 km on either side of the railway. We exclude railways tagged as abandoned or disused.

Importantly, neither gRoads nor OSM datasets provide true and comprehensive temporal information (gRoads not at all); as such both datasats were used in their most up-to-date version in all time periods considered.

### Navigable waterways

Like roads, coastlines and navigable rivers act as conduits for people to access nature. While all coastlines are theoretically navigable, for the purposes of the human footprint we only considered coasts^96^ as navigable for 80 km either direction of signs of a human settlement, which were mapped as a night lights signal with a DN^72^ greater than 6 within 4 km of the coast. We chose 80 km as an approximation of the distance a vessel can travel and return during daylight hours. As new settlements can arise to make new sections of coast navigable, coastal layers were generated for the years 2000, 2005, 2010, and 2013.

Large lakes can act essentially as inland seas, with their coasts frequently plied by trade and harvest vessels. Based on their size and visually identified shipping traffic and shore side settlements, we treated the great lakes of North America, Lake Nicaragua, Lake Titicaca in South America, Lakes Onega and Peipus in Russia, Lakes Balkash and Issyk Kul in Kazakhstan, and Lakes Victoria, Tanganyika and Malawi in Africa as we did navigable marine coasts.

Rivers were considered as navigable if their depth was greater than 2 m and there were signs of night-time lights (DN>=6) within 4 km of their banks, or if contiguous with a navigable coast or large inland lake, and then for a distance of 80 km or until stream depth is likely to prevent boat traffic. To map rivers and their depth we used the hydrosheds (hydrological data and maps based on shuttle elevation derivatives at multiple scales)^87^ dataset on stream discharge, and the following formulae^97,98^:

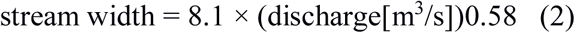

and

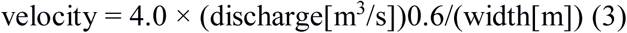

and

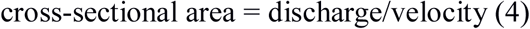

and

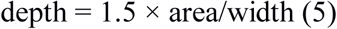

Assuming second order parabola as channel shape.

Navigable rivers layers were created for the years 2000, 2005, 2010, and 2013, and combined with the navigable coasts and inland seas layers for the same years to create the final navigable waterways layers. The access pressure from navigable water bodies were awarded a score of 4 adjacent to the water body, decaying exponentially out to 15 km.

#### Defining low-pressure areas and wilderness

We defined intact areas with low human pressure as a human footprint value of <4, and the areas of high human pressure, or ‘damaged’ areas, as ≥4. This value of ≥4 equates to a human pressure score equal to pasture lands, representing a reasonable approximation of when anthropogenic land conversion has occurred to an extent that the land can be considered human-dominated and no longer ‘natural’. This threshold, which is considered significant at the landscape level^25^, is also the point where species are far more likely to be threatened by habitat loss^12^.

Within the intact state, we defined areas that are pressure-free, or ‘wilderness’, as a human footprint value of <1 following previous global wilderness assessments^10^. We defined wilderness because it increasingly holds special importance in global policy dialogue, as they contain the highest densities of Earth’s biomass, remaining intact mega-faunal assemblages, provide life-supporting ecosystem services, act as controls against which to measure planetary health, provide the last strongholds for many of the world’s languages and have spiritual and cultural value for many of the world’s people of many religions^7,8,99,100^.

#### Units of analysis

Biomes and ecoregions are ecologically distinct geographical units that reflect the distributions of a broad range of fauna and flora across the entire planet ^27^. These entities are now critical for policy and decision makers, being considered core units of reporting in global treaties, and as such can direct legislation, management and conservation efforts towards crisis locations and ecosystems^6,30,31,101,102^. We use biomes and ecoregions described by Olson and colleagues in 2001^27^ to define terrestrial biomes and ecoregions, excluding Lakes and Rock and Ice. We excluded ecoregions that either fell within the Lakes, Rock and Ice biomes or were not covered by the human footprint. World borders were described by Sandvik 2009^103^, both datasets are freely downloadable.

#### Assessing human footprint change

We calculated transitions in levels of human pressure by first assessing human footprint scores for the year 2000, then identified pixels that had changed to a different intensity through to the year 2013. We assume that once a pixel has moved from a score of 0 (a wilderness state), it cannot return to this condition as by definition, once transformed an area is no longer wilderness^8,104^. Therefore, any pixel that was <1 in 2013, but greater than 1 in any other year was given a value of 1 so that it is considered intact land rather than wilderness. All other comparisons directly report changes between 2000 and 2013, including positive changes when a pixel has a lower human footprint value in the year 2013 than it did in the year 2000. We assess both total area and proportional losses, as smaller losses in smaller units may potentially more significant to those unique assemblages as large ones ^105^. In addition to calculating the overall state of biomes and ecoregions for the years 2000 and 2013, we calculated the state for each time period in the human footprint dataset (2000, 2005, 2010 and 2013). All spatial analyses were carried out using ArcMap 10.5^106^. We report on values rounded to the nearest ten throughout for readability. For all values see Supplemental 1.

## Acknowledgments

B.W. and M.W. were supported by an Australian Government Research Training Program Scholarship. The work was funded by the NASA Biodiversity and Ecological Forecasting Program under the 2016 ECO4CAST solicitation through grant NNX17AG51G.

## Author Contributions

J.W conceived the idea and B.W. and J.W. designed the research. B.W. carried out the analysis and led the writing of the manuscript. O.V and S.A created the updated human footprint maps with the support of J.E, S.G, A.H, P.J, R.P, S.R.B, C.S, and A.V. All authors contributed to and edited the manuscript.

## Supplementary material

Supplemental 1 – Excel sheets detailing the area in each state, and the area that transitioned between each state at the global, biome, ecoregional, and national scales.

Supplemental 2 – Technical validation for the human footprint

Table S1 – Summary of the data and methodology used to create the human footprint maps for the years 2000, 2005, 2010 and 2013

## Data availability

The updated human footprint maps, and all the code for generating them are freely downloadable from https://github.com/scabecks/humanfootprint_2000-2013. All other geographic layers used to carry out this analysis are available online from the reference sources.

## References

1. Ellis, E.C., and Ramankutty, N. (2008). Putting people in the map: anthropogenic biomes of the world. Front. Ecol. Environ. 6, 439–447.

2. Morales-Hidalgo, D., Oswalt, S.N., and Somanathan, E. (2015). Status and trends in global primary forest, protected areas, and areas designated for conservation of biodiversity from the Global Forest Resources Assessment 2015. For. Ecol. Manage. 352, 68–77.

3. Hazlitt, S.L., Martin, T.G., Sampson, L., and Arcese, P. (2010). The effects of including marine ecological values in terrestrial reserve planning for a forest-nesting seabird. Biol. Conserv. 143, 1299–1303.

4. Bonan, G.B. (2008). Forests and climate change: forcings, feedbacks, and the climate benefits of forests. Science. 320, 1444–1449.

5. Sheil, D., and Murdiyarso, D. (2009). How forests attract rain: an examination of a new hypothesis. Bioscience 59, 341–347.

6. Dinerstein, E., Vynne, C., Sala, E., Joshi, A.R., Fernando, S., Lovejoy, T.E., Mayorga, J., Olson, D., Asner, G.P., and Baillie, J.E.M. (2019). A Global Deal For Nature: Guiding principles, milestones, and targets. Sci. Adv. 5, eaaw2869.

7. Lovejoy, T.E. (2016). Conservation biology: the importance of wilderness. Curr. Biol. 26, R1235–R1237.

8. Watson, J.E.M., Venter, O., Lee, J., Jones, K.R., Robinson, J.G., Possingham, H.P., and Allan, J.R. (2018). Protect the last of the wild. Nature 563, 27–30.

9. Watson, J.E.M., and Venter, O. (2019). Mapping the continuum of humanity’s footprint on land. OneEarth 1, 175–180.

10. Allan, J.R., Venter, O., and Watson, J.E.M. (2017). Temporally inter-comparable maps of terrestrial wilderness and the Last of the Wild. Sci. data 4, 170187.

11. Jones, K.R., Klein, C.J., Halpern, B.S., Venter, O., Grantham, H., Kuempel, C.D., Shumway, N., Friedlander, A.M., Possingham, H.P., and Watson, J.E.M. (2018). The location and protection status of Earth’s diminishing marine wilderness. Curr. Biol. 28, 2506–2512.

12. Di Marco, M., Venter, O., Possingham, H.P., and Watson, J.E.M. (2018). Changes in human footprint drive changes in species extinction risk. Nat. Commun. 9, 4621.

13. Beyer, H.L., Venter, O., Grantham, H.S., and Watson, J.E.M. (2019). Substantial losses in ecoregion intactness highlight urgency of globally coordinated action. Conserv. Lett. e12592.

14. Jones, K.R., Venter, O., Fuller, R.A., Allan, J.R., Maxwell, S.L., Negret, P.J., and Watson, J.E.M. (2018). One-third of global protected land is under intense human pressure. Science. 360, 788–791.

15. Allan, J.R., Watson, J.E.M., Di, M.M., O’Bryan, C.J., Possingham, H.P., Atkinson, S.C., and Venter, O. (2019). Hotspots of human impact on threatened terrestrial vertebrates. PLoS Biol. 17, e3000158.

16. Tucker, M.A., Böhning-Gaese, K., Fagan, W.F., Fryxell, J.M., Van Moorter, B., Alberts, S.C., Ali, A.H., Allen, A.M., Attias, N., and Avgar, T. (2018). Moving in the Anthropocene: Global reductions in terrestrial mammalian movements. Science. 359, 466–469.

17. Allan, J.R., Venter, O., Maxwell, S., Bertzky, B., Jones, K., Shi, Y., and Watson, J.E.M. (2017). Recent increases in human pressure and forest loss threaten many Natural World Heritage Sites. Biol. Conserv. 206, 47–55.

18. Geldmann, J., Joppa, L.N., and Burgess, N.D. (2014). Mapping change in human pressure globally on land and within protected areas. Conserv. Biol. 28, 1604–1616.

19. Watson, J.E.M., and Venter, O. (2017). A global plan for nature conservation. Nature 550, 48–49.

20. Tulloch, V.J.D., Tulloch, A.I.T., Visconti, P., Halpern, B.S., Watson, J.E.M., Evans, M.C., Auerbach, N.A., Barnes, M., Beger, M., and Chadès, I. (2015). Why do we map threats? Linking threat mapping with actions to make better conservation decisions. Front. Ecol. Environ. 13, 91–99.

21. Vörösmarty, C.J., McIntyre, P.B., Gessner, M.O., Dudgeon, D., Prusevich, A., Green, P., Glidden, S., Bunn, S.E., Sullivan, C.A., and Liermann, C.R. (2010). Global threats to human water security and river biodiversity. Nature 467, 555–561.

22. Allan, J.R., Grossmann, F., Craig, R., Nelson, A., Maina, J., Flower, K., Bampton, J., ste Deffontaines, J.-B., Miguel, C., and Araquechande, B. (2017). Patterns of forest loss in one of Africa’s last remaining wilderness areas: Niassa National Reserve (Northern Mozambique). Parks 23, 39–50.

23. Venter, O., Sanderson, E.W., Magrach, A., Allan, J.R., Beher, J., Jones, K.R., Possingham, H.P., Laurance, W.F., Wood, P., and Fekete, B.M. (2016). Global terrestrial Human Footprint maps for 1993 and 2009. Sci. data 3, sdata201667.

24. Sanderson, E.W., Jaiteh, M., Levy, M.A., Redford, K.H., Wannebo, A. V, and Woolmer, G. (2002). The human footprint and the last of the wild: the human footprint is a global map of human influence on the land surface, which suggests that human beings are stewards of nature, whether we like it or not. AIBS Bull. 52, 891–904.

25. Watson, J.E.M., Jones, K.R., Fuller, R.A., Marco, M. Di, Segan, D.B., Butchart, S.H.M., Allan, J.R., McDonald-Madden, E., and Venter, O. (2016). Persistent disparities between recent rates of habitat conversion and protection and implications for future global conservation targets. Conserv. Lett. 9, 413–421.

26. Crooks, K.R., Burdett, C.L., Theobald, D.M., King, S.R.B., Di Marco, M., Rondinini, C., and Boitani, L. (2017). Quantification of habitat fragmentation reveals extinction risk in terrestrial mammals. Proc. Natl. Acad. Sci. 114, 7635–7640.

27. Olson, D.M.D., Dinerstein, E., Wikramanayake, E.D., Burgess, N.D., Powell, G.V.N., Underwood, E.C., D’amico, J.A., Itoua, I., Strand, H.E., and Morrison, J.C. (2001). Terrestrial Ecoregions of the World: A New Map of Life on Earth: A new global map of terrestrial ecoregions provides an innovative tool for conserving biodiversity. Bioscience 51, 933–938.

28. Newbold, T., Hudson, L.N., Arnell, A.P., Contu, S., De Palma, A., Ferrier, S., Hill, S.L.L., Hoskins, A.J., Lysenko, I., and Phillips, H.R.P. (2016). Has land use pushed terrestrial biodiversity beyond the planetary boundary? A global assessment. Science. 353, 288–291.

29. Steffen, W., Richardson, K., Rockström, J., Cornell, S.E., Fetzer, I., Bennett, E.M., Biggs, R., Carpenter, S.R., De Vries, W., and De Wit, C.A. (2015). Planetary boundaries: Guiding human development on a changing planet. Science. 347, 1259855.

30. Dinerstein, E., Olson, D., Joshi, A., Vynne, C., Burgess, N.D., Wikramanayake, E., Hahn, N., Palminteri, S., Hedao, P., and Noss, R. (2017). An ecoregion-based approach to protecting half the terrestrial realm. Bioscience 67, 534–545.

31. Hoekstra, J.M., Boucher, T.M., Ricketts, T.H., and Roberts, C. (2005). Confronting a biome crisis: global disparities of habitat loss and protection. Ecol. Lett. 8, 23–29.

32. Hannah, L., Carr, J.L., and Lankerani, A. (1995). Human disturbance and natural habitat: a biome level analysis of a global data set. Biodivers. Conserv. 4, 128–155.

33. Tulloch, A.I.T., Gordon, A., Runge, C.A., and Rhodes, J.R. (2019). Integrating spatially realistic infrastructure impacts into conservation planning to inform strategic environmental assessment. Conserv. Lett., e12648.

34. Gaston, K.J., Visser, M.E., and Hölker, F. (2015). The biological impacts of artificial light at night: the research challenge.

35. Hampton, S.E., Strasser, C.A., Tewksbury, J.J., Gram, W.K., Budden, A.E., Batcheller, A.L., Duke, C.S., and Porter, J.H. (2013). Big data and the future of ecology. Front. Ecol. Environ. 11, 156–162.

36. Carr, D.L. (2004). Proximate population factors and deforestation in tropical agricultural frontiers. Popul. Environ. 25, 585–612.

37. Watson, J.E.M., Evans, T., Venter, O., Williams, B., Tulloch, A., Stewart, C., Thompson, I., Ray, J.C., Murray, K., Salazar, A., et al. (2018). The exceptional value of intact forest ecosystems. Nat. Ecol. Evol. 2, 599–610.

38. Fleischner, T.L. (1994). Ecological costs of livestock grazing in western North America. Conserv. Biol. 8, 629–644.

39. Newbold, T., Hudson, L.N., Hill, S.L.L., Contu, S., Lysenko, I., Senior, R.A., Börger, L., Bennett, D.J., Choimes, A., and Collen, B. (2015). Global effects of land use on local terrestrial biodiversity. Nature 520, 45.

40. Miraldo, A., Li, S., Borregaard, M.K., Flórez-Rodríguez, A., Gopalakrishnan, S., Rizvanovic, M., Wang, Z., Rahbek, C., Marske, K.A., and Nogués-Bravo, D. (2016). An Anthropocene map of genetic diversity. Science. 353, 1532–1535.

41. Maxwell, S.L., Evans, T., Watson, J.E.M., Morel, A., Grantham, H., Duncan, A., Harris, N., Potapov, P., Runting, R.K., and Venter, O. (2019). Degradation and forgone removals increase the carbon impact of intact forest loss by 626. Sci. Adv. 5, eaax2546.

42. Vargas, L.E.P., Laurance, W.F., Clements, G.R., and Edwards, W. (2015). The impacts of oil palm agriculture on Colombia’s biodiversity: what we know and still need to know. Trop. Conserv. Sci. 8, 828–845.

43. Williams, B.A., Grantham, H.S., Watson, J.E.M., Alvarez, S.J., Simmonds, J.S., Rogéliz, C.A., Da Silva, M.A., Forero-Medina, G., Etter, A., Nogales, J., et al. (2020). Minimising the loss of biodiversity and ecosystem services in an intact landscape under risk of rapid agricultural development. Environ. Res. Lett. 15, 14001.

44. Australian Government (2015). Our north, our future: White paper on developing northern Australia Available at: https://www.industry.gov.au/data-and-publications/our-north-our-future-white-paper-on-developing-northern-australia.

45. Strassburg, B.B.N., Brooks, T., Feltran-Barbieri, R., Iribarrem, A., Crouzeilles, R., Loyola, R., Latawiec, A.E., Oliveira Filho, F.J.B., Scaramuzza, C.A. de M., and Scarano, F.R. (2017). Moment of truth for the Cerrado hotspot. Nat. Ecol. Evol. 1, 99.

46. Monteiro, L.M., Brum, F.T., Pressey, R.L., Morellato, L.P.C., Soares-Filho, B., Lima-Ribeiro, M.S.,, and Loyola, R. (2018). Evaluating the impact of future actions in minimizing vegetation loss from land conversion in the Brazilian Cerrado under climate change. Biodivers. Conserv. 29, 1–22.

47. Pressey, R.L., and Taffs, K.H. (2001). Scheduling conservation action in production landscapes: priority areas in western New South Wales defined by irreplaceability and vulnerability to vegetation loss. Biol. Conserv. 100, 355–376.

48. Wilson, K., Pressey, R.L., Newton, A., Burgman, M., Possingham, H., and Weston, C. (2005). Measuring and incorporating vulnerability into conservation planning. Environ. Manage. 35, 527–543.

49. Lenton, T.M., Rockström, J., Gaffney, O., Rahmstorf, S., Richardson, K., Steffen, W., and Schellnhuber, H.J. (2019). Climate tipping points—too risky to bet against. Nature 575, 592–595.

50. Di Marco, M., Ferrier, S., Harwood, T.D., Hoskins, A.J., and Watson, J.E.M. (2019). Wilderness areas halve the extinction risk of terrestrial biodiversity. Nature 573, 582–585.

51. IPBES (2019). Global assessment report on biodiversity and ecosystem services of the Intergovernmental Science-Policy Platform on Biodiversity and Ecosystem Services (Bonn, Germany) Available at: https://www.ipbes.net/global-assessment-report-biodiversity-ecosystem-services.

52. Visseren-Hamakers, I.J., (2015). Integrative environmental governance: enhancing governance in the era of synergies. Curr. Opin. Environ. Sustain. 14, 136–143.

53. Wunder, S. (2007). The efficiency of payments for environmental services in tropical conservation. Conserv. Biol. 21, 48–58.

54. Maron, M., Simmonds, J.S., and Watson, J.E.M. (2018). Bold nature retention targets are essential for the global environment agenda. Nat. Ecol. Evol. 2, 1194.

55. Watson, J.E.M., Keith, D.A., Strassburg, B.B.N., Venter, O., Williams, B., and Nicholson, E. (2020). Set a global target for ecosystems. Nature 578, 360–362.

56. Di Marco, M., Harwood, T.D., Hoskins, A.J., Ware, C., Hill, S.L.L., and Ferrier, S. (2019). Projecting impacts of global climate and land-use scenarios on plant biodiversity using compositional-turnover modelling. Glob. Chang. Biol. 25, 2763–2778.

57. Van Ittersum, M.K., Van Bussel, L.G.J., Wolf, J., Grassini, P., Van Wart, J., Guilpart, N., Claessens, L., de Groot, H., Wiebe, K., and Mason-D’Croz, D. (2016). Can sub-Saharan Africa feed itself? Proc. Natl. Acad. Sci. 113, 14964–14969.

58. Shiferaw, B., Negassa, A., Koo, J., Wood, J., Sonder, K., Braun, J.A., and Payne, T. (2011). Future of wheat production in Sub-Saharan Africa: analyses of the expanding gap between supply and demand and economic profitability of domestic production. In Increasing Agricultural Productivity & Enhancing Food Security in Africa: New Challenges and Opportunities (Africa Hall, UNECA, Addis Ababa, Ethiopia: International Food Policy Research Institute (IFPRI)).

59. Margono, B.A., Potapov, P. V, Turubanova, S., Stolle, F., and Hansen, M.C. (2014). Primary forest cover loss in Indonesia over 2000-2012. Nat. Clim. Chang. 4, 730.

60. Australian government About Papua New Guinea. Available at: https://web.archive.org/web/20110518125558/http://www.ausaid.gov.au/country/png/png_intro.cfm.

61. Nelson, P.N., Gabriel, J., Filer, C., Banabas, M., Sayer, J.A., Curry, G.N., Koczberski, G., and Venter, O. (2014). Oil palm and deforestation in Papua New Guinea. Conserv. Lett. 7, 188–195.

62. Austin, K.G., Mosnier, A., Pirker, J., McCallum, I., Fritz, S., and Kasibhatla, P.S. (2017). Shifting patterns of oil palm driven deforestation in Indonesia and implications for zero-deforestation commitments. Land use policy 69, 41–48.

63. Lenzen, M., Moran, D., Kanemoto, K., Foran, B., Lobefaro, L., and Geschke, A. (2012). International trade drives biodiversity threats in developing nations. Nature 486, 109.

64. Costanza, R., Daly, L., Fioramonti, L., Giovannini, E., Kubiszewski, I., Mortensen, L.F., Pickett, K.E., Ragnarsdottir, K.V., De Vogli, R., and Wilkinson, R. (2016). Modelling and measuring sustainable wellbeing in connection with the UN Sustainable Development Goals. Ecol. Econ. 130, 350–355.

65. Secretariat of the Convention on Biological Diversity (2020). Zero Draft of the Post-2020 Global Biodiversity Framework Available at: https://www.cbd.int/article/2020-01-10-19-02-38.

66. OS Geo Project (2017). GRASS GIS 7.2.2. Available at: https://grass.osgeo.org/news/68/15/GRASS-GIS-7-2-2-released/.

67. Tratalos, J., Fuller, R.A., Warren, P.H., Davies, R.G., and Gaston, K.J. (2007). Urban form, biodiversity potential and ecosystem services. Landsc. Urban Plan. 83, 308–317.

68. Aronson, M.F.J., La Sorte, F.A., Nilon, C.H., Katti, M., Goddard, M.A., Lepczyk, C.A., Warren, P.S., Williams, N.S.G., Cilliers, S., and Clarkson, B. (2014). A global analysis of the impacts of urbanization on bird and plant diversity reveals key anthropogenic drivers. Proc. R. Soc. B Biol. Sci. 281, 20133330.

69. Butchart, S.H.M., Walpole, M., Collen, B., Van Strien, A., Scharlemann, J.P.W., Almond, R.E.A., Baillie, J.E.M., Bomhard, B., Brown, C., and Bruno, J. (2010). Global biodiversity: indicators of recent declines. Science. 328, 1164–1168.

70. Chamberlain, D.E., Cannon, A.R., Toms, M.P., Leech, D.I., Hatchwell, B.J., and Gaston, K.J. (2009). Avian productivity in urban landscapes: a review and meta-analysis. Ibis (Lond. 1859). 151, 1–18.

71. Elvidge, C.D., Hsu, F.-C., Baugh, K.E., and Ghosh, T. (2014). National trends in satellite-observed lighting. Glob. urban Monit. Assess. through earth Obs. 23, 97–118.

72. Elvidge, C.D., Imhoff, M.L., Baugh, K.E., Hobson, V.R., Nelson, I., Safran, J., Dietz, J.B., and Tuttle, B.T. (2001). Night-time lights of the world: 1994-1995. ISPRS J. Photogramm. Remote Sens. 56, 81–99.

73. Small, C., Elvidge, C.D., Balk, D., and Montgomery, M. (2011). Spatial scaling of stable night lights. Remote Sens. Environ. 115, 269–280.

74. Elvidge, C.D., Ziskin, D., Baugh, K.E., Tuttle, B.T., Ghosh, T., Pack, D.W., Erwin, E.H., and Zhizhin, M. (2009). A fifteen year record of global natural gas flaring derived from satellite data. Energies 2, 595–622.

75. Elvidge, C.D., Baugh, K.E., Zhizhin, M., and Hsu, F.-C. (2013). Why VIIRS data are superior to DMSP for mapping nighttime lights. Proc. Asia-Pacific Adv. Netw. 35.

76. Brashares, J.S., Arcese, P., and Sam, M.K. (2001). Human demography and reserve size predict wildlife extinction in West Africa. Proc. R. Soc. London. Ser. B Biol. Sci. 268, 2473–2478.

77. Miller, G.H., Fogel, M.L., Magee, J.W., Gagan, M.K., Clarke, S.J., and Johnson, B.J. (2005). Ecosystem collapse in Pleistocene Australia and a human role in megafaunal extinction. Science. 309, 287–290.

78. Burney, D.A., and Flannery, T.F. (2005). Fifty millennia of catastrophic extinctions after human contact. Trends Ecol. Evol. 20, 395–401.

79. CIESIN, and SEDAC (2017). Gridded Population of the World Version 4. Cent. Int. Earth Sci. Inf. Netw., 1–21. Available at: http://sedac.ciesin.columbia.edu/data/set/gpw-v4-population-density [Accessed September 11, 2017].

80. NOAA (2013). Version 4 DMSP-OLS Nighttime Lights Time Series. Available at: https://ngdc.noaa.gov/eog/dmsp/downloadV4composites.html#AVSLCFC [Accessed March 17, 2020].

81. ESA (2017). 300 m annual global land cover time series from 1992 to 2015. Available at: http://maps.elie.ucl.ac.be/CCI/viewer/ [Accessed July 13, 2017].

82. Fischer, J., Brosi, B., Daily, G.C., Ehrlich, P.R., Goldman, R., Goldstein, J., Lindenmayer, D.B., Manning, A.D., Mooney, H.A., and Pejchar, L. (2008). Should agricultural policies encourage land sparing or wildlife-friendly farming? Front. Ecol. Environ. 6, 380–385.

83. Ramankutty, N., Evan, A.T., Monfreda, C., and Foley, J.A. (2008). Farming the planet: 1. Geographic distribution of global agricultural lands in the year 2000. Global Biogeochem. Cycles 22, GB1003.

84. Luck, G.W., and Daily, G.C. (2003). Tropical countryside bird assemblages: richness, composition, and foraging differ by landscape context. Ecol. Appl. 13, 235–247.

85. OpenStreetMap Contributors (2017). Planet OSM. Available at: https://planet.osm.org [Accessed May 29, 2017].

86. Herold, M., Mayaux, P., Woodcock, C.E., Baccini, A., and Schmullius, C. (2008). Some challenges in global land cover mapping: An assessment of agreement and accuracy in existing 1 km datasets. Remote Sens. Environ. 112, 2538–2556.

87. Lehner, B., Verdin, K., and Jarvis, A. (2008). New global hydrography derived from spaceborne elevation data. Eos, Trans. Am. Geophys. Union 89, 93–94.

88. Kauffman, J.B., and Krueger, W.C. (1984). Livestock impacts on riparian ecosystems and streamside management implications... a review. Rangel. Ecol. Manag. Range Manag. Arch. 37, 430–438.

89. Trombulak, S.C., and Frissell, C.A. (2000). Review of ecological effects of roads on terrestrial and aquatic communities. Conserv. Biol. 14, 18–30.

90. Woodroffe, R., and Ginsberg, J.R. (1998). Edge effects and the extinction of populations inside protected areas. Science. 280, 2126–2128.

91. Laurance, W.F., Goosem, M., and Laurance, S.G.W. (2009). Impacts of roads and linear clearings on tropical forests. Trends Ecol. Evol. 24, 659–669.

92. Adeney, J.M., Christensen Jr, N.L., and Pimm, S.L. (2009). Reserves protect against deforestation fires in the Amazon. PLoS One 4, e5014.

93. Forman, R.T.T., and Alexander, L.E. (1998). Roads and their major ecological effects. Annu. Rev. Ecol. Syst. 29, 207–231.

94. OpenStreetMap, and OpenStreetMap Contributors (2020). OpenStreetMap. Available at: https://www.openstreetmap.org/about [Accessed March 18, 2020].

95. Center for International Earth Science Information Network (2010). Global Roads Open Access Data Set (gROADS), v1 (1980 – 2010). NASA Socioecon. Data Appl. Cent. Available at: https://sedac.ciesin.columbia.edu/data/set/groads-global-roads-open-access-v1.

96. National Imagery and Mapping Agency (1997). National Imagery and Mapping Agency. Vector Map Level 0. Available at: https://earth-info.nga.mil/publications/vmap0.html.

97. Bjerklie, D.M., Dingman, S.L., Vorosmarty, C.J., Bolster, C.H., and Congalton, R.G. (2003). Evaluating the potential for measuring river discharge from space. J. Hydrol. 278, 17–38.

98. Bjerklie, D.M., Moller, D., Smith, L.C., and Dingman, S.L. (2005). Estimating discharge in rivers using remotely sensed hydraulic information. J. Hydrol. 309, 191–209.

99. Mittermeier, R.A., Mittermeier, C.G., Brooks, T.M., Pilgrim, J.D., Konstant, W.R., Da Fonseca, G.A.B., and Kormos, C. (2003). Wilderness and biodiversity conservation. Proc. Natl. Acad. Sci. 100, 10309–10313.

100. Watson, J.E.M., Shanahan, D.F., Di Marco, M., Allan, J., Laurance, W.F., Sanderson, E.W., Mackey, B., and Venter, O. (2016). Catastrophic declines in wilderness areas undermine global environment targets. Curr. Biol. 26, 2929–2934.

101. Forero-Medina, G., and Joppa, L. (2010). Representation of global and national conservation priorities by Colombia’s protected area network. PLoS One 5, e13210.

102. Rodríguez, J.P., Rodríguez-Clark, K.M., Baillie, J.E.M., Ash, N., Benson, J., Boucher, T., Brown, C., Burgess, N.D., Collen, B.E.N., and Jennings, M. (2011). Establishing IUCN red list criteria for threatened ecosystems. Conserv. Biol. 25, 21–29.

103. Sandvik, B. (2009). World Borders Dataset. Available at: thematicmapping.org.

104. Crouzeilles, R., Curran, M., Ferreira, M.S., Lindenmayer, D.B., Grelle, C.E. V, and Benayas, J.M.R. (2016). A global meta-analysis on the ecological drivers of forest restoration success. Nat. Commun. 7, 1–8.

105. He, F., and Hubbell, S.P. (2011). Species–area relationships always overestimate extinction rates from habitat loss. Nature 473, 368–371.

106. ESRI (2017). ArcGIS Release 10.5.1. Redlands, CA.

