## Supplemental 2 for "Change in terrestrial human footprint drives continued loss of intact ecosystems"

Supplemental 2 - Technical validation for the human footprint

High resolution images were used to visually interpret human pressures in 3460 × 1 km^2^ sample plots randomly located across the Earth’s non-Antarctic land areas. Images for these plots were obtained from World Imagery^1^, which provides one meter or better satellite and aerial imagery in many parts of the world and lower resolution satellite imagery worldwide. The map features 0.3m resolution imagery across the continental United States and parts of Western Europe, as well as many parts of the world, with concentrations in South America, Eastern Europe, India, Japan, the Middle East and Northern Africa, Southern Africa, Australia, and New Zealand. The imagery used for the validation plots had a median resolution of 0.5 meters and a median acquisition year of 2010. Therefore, only the 2010 map underwent validation.

For the visual interpretation, the extent of built environments, crop lands, pasture lands, roads, human settlements, infrastructures and navigable waterways, were recorded using a standard key for identifying these features (See Supplementary Appendix 1 of Venter et al. 2016^2^). Shape, size, texture and colour of features in the imagery were important characteristics for identifying human pressures on the environment. Interpretations were also marked as ‘certain’ or ‘uncertain’, and the year and resolution of the interpreted image was recorded. The 346 ‘uncertain’ plots were discarded, leaving 3114 validation plots. In general, plots were classified as ‘uncertain’ for two reasons; either because cloud cover obscured the image, or because only medium resolution (15 m) imagery was available for the plot, preventing accurate interpretation of the image. The human footprint score for each plot was determined in ArcGIS, and the visual and Human Footprint scores were then normalized to a 0–1 scale. As we only retained plots for which visual interpretations of the images were determined to be ‘certain’, we consider the visual score to be the true state of in-situ pressures for the plots.

Two statistics were used to determine Human Footprint performance, root mean squared error (RMSE)^3^ and the Cohen kappa statistic of agreement^4^. The RMSE is a dimensioned (expresses average error in the units of variable of interest) error metric for numerical predictions, and tends to heavily punish large errors. The Kappa statistic expresses the agreement between two categorical datasets corrected for the expected agreement, which is based on a random allocation given the relative class sizes. When calculating the kappa statistic, the Human Footprint score was considered as a match to the visual

score if they were within 20% of one another on the 0–1 scale.

There is strong agreement between the Human Footprint measure of pressure and pressures scored by visual interpretation of high resolution imagery. The RMSE for the 3114 validation plots was 0.116 on the normalized 0-1 scale. The Kappa statistic was 0.806 (P<0.01), also indicating strong agreement between the Human Footprint and the validation dataset. Of the 3114 1 km^2^ validation plots, the Human Footprint scored 143 of them 20% higher than the visual score and 140 of them 20% lower, indicating a balance between overestimation and underestimation of pressures. The remaining 2831 plots (90.9%) were within 20% agreement. The Kappa statistic measure of agreement is sensitive to the threshold used to consider plots as a ‘match’. If we apply a more stringent threshold for agreement of within 15% of one another, the Kappa statistic falls to 0.645 (moderate agreement), and if we apply a less stringent threshold of within 25%, the Kappa statistic

increases to 0.906 (almost perfect). While agreement is generally strong, there is some geographic variation in the RMSE results comparing the Human Footprint scores and those derived from visual interpretation. By calculating RMSE for all biomes that contain at least 100 of the 3114 sample plots, we found that agreement was strongest in Montane grasslands biome and the Temperate grasslands, savannah and shrubs biome. Agreement was weakest in the Tropical and subtropical moist broadleaf forests and the Deserts and xeric shrublands biomes.

Table S2A - Root Mean Square Errors results comparing the Human Footprint scores with 3114 validation plots globally, and for biomes with at least 100 plots within them.

| **Region** | **RMSE** |
| --- | --- |
| RMSE Global | 0.115551 |
| RMSE Boreal | 0.110667 |
| RMSE Deserts and xeric shrublands | 0.124895 |
| RMSE Montane grasslands | 0.093302 |
| RMSE Temperate broadleaf and mixed forests | 0.112171 |
| RMSE Temperate grasslands, savannahs, and shrublands | 0.108985 |
| RMSE Tropical and subtropical grasslands, savannahs, and shrublands | 0.109195 |
| RMSE Tropical and subtropical moist broadleaf forests | 0.116537 |
| RMSE Tundra | 0.112789 |

**References**

1.	ESRI (2016). World Imagery. Available at: http://services.arcgisonline.com/ArcGIS/rest/services/World_Imagery/MapServer.

2.	Venter, O., Sanderson, E.W., Magrach, A., Allan, J.R., Beher, J., Jones, K.R., Possingham, H.P., Laurance, W.F., Wood, P., and Fekete, B.M. (2016). Global terrestrial Human Footprint maps for 1993 and 2009. Sci. data *3*, sdata201667.

3.	Willmott, C.J., and Matsuura, K. (2005). Advantages of the mean absolute error (MAE) over the root mean square error (RMSE) in assessing average model performance. Clim. Res. *30*, 79–82.

4.	Viera, A.J., and Garrett, J.M. (2005). Understanding interobserver agreement: the kappa statistic. Fam med *37*, 360–363.
