## Supplementary material for "Change in terrestrial human footprint drives continued loss of intact ecosystems": Table S1

Table S1 - Summary of the data and methodology used to create the human footprint maps for the years 2000, 2005, 2010 and 2013

| **Description** | **Source** | **Data used** | **Temporal range** | **Source Resolution** | **Data processing** | **Outputs used in Human Footprint** |
| --- | --- | --- | --- | --- | --- | --- |
| Built environments | 1–3, personal comms for year 2013 coefficients. | Average, stable lights, & cloud free coverages | 2000 – 2013 | 30 arc second, ~1 km at equator | 1) Preprocessing to reclass all digital number (DN) values < 6 to 0  2) Inter-calibrate across range of years 3) Re-project and resample to 1 km raster basemap (Mollweide projection) using a bilinear interpolation methodology 4) Convert to binary map of areas exhibiting a DN equal to or above 20 5) Assign these areas a pressure score of 10. | built_areas2000.tif, built_areas2005.tif, built_areas2010.tif,  built_areas2013.tif |
| Population density | 4 | Gridded Population of the World (GPWv4), density grids | 2000, 2005, 2010, 2015 | 30 arc second, ~1 km at equator | 1) Reproject and resample to 1 km raster basemap (Mollweide projection)  3) Linearly interpolate values for year 2013 from 2010/2015 data 2) Assign pressure score using equation (1) in methods | popden2000.tif, popden2005.tif, popden2010.tif,  popden2013.tif |
| Night-time lights | 1–3 | Average, stable lights, & cloud free coverages | 2000 – 2013 | 30 arc second, ~1 km at equator | 1) Intercalibrate across years. 2) Reproject and resample to 1 km raster basemap (Mollweide projection.) 3) Create 10 equal quantile bins for year 2000. 4) Assign pressure scores to bins from 0–10, for 2000, and using the same DN thresholds for all other years. | nightlights2000.tif, nightlights2005.tif, nightlights2010.tif, nightlights2013.tif |
| Crop lands | 5 | CCI Land cover | 2000 – 2013 | 300 m | 1) Reproject and resample to 1 km raster basemap (using mode resampling) (Mollweide projection) 2) Convert to binary map showing only crop lands (pixels classified as crops – e.g., 10, 11, 12, 20) 3) Reclass all areas already mapped as built to 0. 4) Assign crop lands a pressure score of 7 | crops_meris2000.tif, crops_meris2005.tif, crops_meris2010.tif, crops_meris2013.tif |
| Pasture lands | 6 | M3-Pasture data | 2000 | 5 min, ~10 km at equator | 1) Reproject and resample to 1 km raster basemap (Mollweide projection) 2) Reclass all areas already mapped as built or crop lands to 0 3) Assign pressure score of 4, weighted by percent pasture lands | pastures2000.tif, pastures2005.tif, pastures2010.tif, pastures2013.tif |
| Roads | 4,7 | OSM roads (Planet.osm dated 29 May 2017); gRoads | 2017 | Vector data | 1) Extract features from OSM where the tag ‘highway’ includes values ‘motorway*’, ’primary*’, ‘residential*’, ‘secondary*’, ‘tertiary*’, ‘trunk*’, ‘track*’, or ‘unclassified*’  2) Merge two datasets  3) Reproject and covert to 1 km raster basemap (Mollweide projection) 4) Assign pressure score of 8 to roaded pixels, and 4 to adjacent pixels, exponentially decaying to 0 at a distance of 15 km | roads.tif |
| Railways | 7 | Railways (Planet.osm dated 29 May 2017) | 2017 | Vector data | 1) Extract features from OSM where the tag ‘railway’ does not include values ‘abandoned’ or ‘disused’  2) Reproject and covert to 1 km raster basemap (Mollweide projection) 2) Assign pressure score of 8 to rail pixels | railways.tif |
| Navigable waterways | 8 | HydroSHEDS, stream discharge | No timeframe | 3 arc second, ~100 m at equator | 1) Reproject and covert to 1 km raster basemap (Mollweide projection) 2) Use equations 2, 3, and 4 to determine stream depth 3) Exclude all stream reaches with water depth less than 2 m deep 4) Exclude all reaches not within 80 km of a stream bank which is within 4 km of a pixel with a DN>4, in the years 2000, 2005, 2010, or 2013 5) Add coastlines within 80 km of a coastal bank which is within 4 km of a pixel with a DN>4, in years 2000, 2005, 2010, or 2015 6) Assign pressure score of 4 to adjacent pixels, exponentially decaying to 0 at distance of 15 km | navwaters2000.tif, navwaters2005.tif, navwaters2010.tif, navwaters2013.tif |

**References**

1.	DMSP (2013). Earth Observation Group - Defense Meteorological Satellite Progam, Boulder. Available at: http://ngdc.noaa.gov/eog/dmsp/downloadV4composites.html#AVSLCFC [Accessed July 11, 2017].

2.	Elvidge, C.D., Hsu, F.-C., Baugh, K.E., and Ghosh, T. (2014). National trends in satellite-observed lighting. Glob. urban Monit. Assess. through earth Obs. *23*, 97–118.

3.	Elvidge, C.D., Imhoff, M.L., Baugh, K.E., Hobson, V.R., Nelson, I., Safran, J., Dietz, J.B., and Tuttle, B.T. (2001). Night-time lights of the world: 1994–1995. ISPRS J. Photogramm. Remote Sens. *56*, 81–99.

4.	CIESIN, and SEDAC (2017). Gridded Population of the World Version 4. Cent. Int. Earth Sci. Inf. Netw., 1–21. Available at: http://sedac.ciesin.columbia.edu/data/set/gpw-v4-population-density [Accessed September 11, 2017].

5.	ESA (2017). 300 m annual global land cover time series from 1992 to 2015. Available at: http://maps.elie.ucl.ac.be/CCI/viewer/ [Accessed July 13, 2017].

6.	Ramankutty, N., Evan, A.T., Monfreda, C., and Foley, J.A. (2008). Farming the planet: 1. Geographic distribution of global agricultural lands in the year 2000. Global Biogeochem. Cycles *22*, GB1003.

7.	OpenStreetMap Contributors (2017). Planet OSM. Available at: https://planet.osm.org [Accessed May 29, 2017].

8.	Lehner, B., Verdin, K., and Jarvis, A. (2008). New global hydrography derived from spaceborne elevation data. Eos, Trans. Am. Geophys. Union *89*, 93–94.
